# Autumn larval cold tolerance does not predict the northern range limit of a widespread butterfly species

**DOI:** 10.1101/2020.06.14.151266

**Authors:** Philippe Tremblay, Heath A. MacMillan, Heather M. Kharouba

## Abstract

Climate change is driving range shifts, and a lack of cold tolerance is hypothesized to constrain insect range expansion at poleward latitudes. However, few, if any, studies have tested this hypothesis during autumn when organisms are subjected to sporadic low temperature exposure but may not have become cold tolerant yet. In this study, we integrated organismal thermal tolerance measures into species distribution models for larvae of the Giant Swallowtail butterfly, *Papilio cresphontes*, living at the northern edge of its actively expanding range. Cold hardiness of field-collected larvae was determined using three common metrics of cold-induced physiological thresholds: the supercooling point (SCP), critical thermal minimum (CT_min_), and survival following cold exposure. *P. cresphontes* larvae in autumn have a CT_min_ of 2.14°C, and were determined to be tolerant of chilling. These larvae have a SCP of −6.6°C and can survive prolonged exposure to −2°C. They generally die, however, at temperatures below their SCP (−8°C), suggesting they are chill tolerant or modestly freeze avoidant. Using this information, we examined the importance of low temperatures at a broad scale, by comparing species distribution models of *P. cresphontes* based only on environmental data derived from other sources to models that also included the cold tolerance parameters generated experimentally. Our modelling revealed that growing degree-days and precipitation best predicted the distribution of *P. cresphontes*, while the cold tolerance variables did not explain much variation in habitat suitability. As such, the modelling results were consistent with our experimental results: low temperatures in autumn are unlikely to limit the distribution of *P. cresphontes*. Further investigation into the ecological relevance of the physiological thresholds determined here will help determine how climate limits the distribution of *P. cresphontes*. Understanding the factors that limit species distributions is key to predicting how climate change will drive species range shifts.

## Introduction

Over the past few decades, range shifts due to climate and land use changes have been reported in both plants and animals. Many are experiencing a poleward and/or upward shift in distribution, pushing their northern and upper range limit to higher latitudes and altitudes (Chen et al., 2011; VanDerWal et al., 2013; but see Kerr et al. 2015). However, there has been substantial variation in the degree and direction of range shift across taxonomic groups (Chen et al., 2011; Lenoir & Svenning 2015). For example, some insect taxa (i.e., dragonflies, spiders) have shifted northwards to a much greater extent than mammals and birds, whereas other groups have expanded southwards. This variation in range shifts remains difficult to explain, and thus necessitates a better understanding of the factors influencing species’ distributions (Buckley & Kingsolver, 2012; Sunday et al. 2012; Sgrò et al., 2016).

A climate change-driven range shift in insects is often attributed to the relaxation of a harsher poleward climate since temperature acts as a physiological limitation and as a phenological cue (Logan & Bentz, 1999; Root et al., 2003; Elkinton et al., 2008). For those species with northern range edges associated with temperature clines, climatic warming has frequently resulted in the colonization of suitable areas in more northern latitudes (Parmesan, 1996; Parmesan & Yohe, 2003). For example, the northern range expansion of the deer fly (*Lipoptena cervi*) in Finland is due to warmer temperatures during the summer (Härkönen et al., 2010). However, for many other species, it remains unclear how climate change has led to range shifts (Chen et al., 2011; Sunday et al., 2012).

Low temperatures throughout the year can constrain the ability of insects to persist at northern range limits as they can influence any life cycle stage (Ungerer et al., 1999). Overwinter survival has been shown to be a key factor limiting the ranges of insects (Crozier, 2003). For example, the northern range edges of the Southern pine beetle (*Dendroctonus frontalis*) and Sachem skipper butterfly (*Atalopedes campestris*) are limited by temperatures below −16°C (Lesk et al., 2017) and the −4°C January isotherm (Crozier, 2003), respectively. Overwintering insects can adopt one of multiple physiological strategies for surviving winter (Williams et al., 2015); some insects can survive internal ice formation and are termed freeze tolerant, some physiologically supress the temperature at which their body fluids spontaneously freeze (the supercooling point; SCP) and are termed freeze avoiding, while others remain susceptible to chilling (termed chill susceptible), and seek relatively warm microhabitats to overwinter (Sinclair, 1999; Denlinger & Lee, 2010; Overgaard & MacMillan, 2017).

While winter represents a considerable challenge (Robinet & Roques, 2010; Williams et al., 2015), insects are also likely to be particularly vulnerable to low temperatures during autumn. Autumn is when most insects are acquiring cold tolerance through physiological adjustments (acclimatization) and have therefore not reached their peak cold hardiness (i.e., the capacity to tolerate intensity and duration of cold exposure; Tauber et al., 1986). For example, monarch butterflies, *Danaus plexippus*, that were not yet acclimated had a mortality rate of 60% when exposed to low temperatures in September and October compared to 20% for acclimated ones (Larsen & Lee, 1994). There is also a much higher frequency of cold temperature events in autumn than in the summer (Danks, 1978). Yet, no study, to our knowledge, has considered low temperatures during autumn as a limiting factor on the geographic distributions of insects.

Regardless of season, cold exposure can cause insects to cross physiological thresholds leading to sublethal effects. At a species-specific low temperature, most insects lose the ability for coordinated movement (a critical thermal minimum; CT_min_) after which they enter a state of complete neuromuscular silence termed chill coma (Gibert & Huey, 2001; MacMillan & Sinclair, 2011; Oyen & Dillon, 2018). Many insects can recover from this state with no immediate evidence of injury; but prolonged or severe cold exposure can cause behavioural defects, or slow or halt development (Asahina, 1970; Rojas & Leopold, 1996; Overgaard & MacMillan, 2017). Chill susceptible insects suffer from a loss of homeostasis at low temperatures well above their SCP, while those more tolerant to chilling survive such exposures (Overgaard & MacMillan, 2017). While freeze tolerant insects can survive freezing of the extracellular fluid at their SCP, they can suffer from ice-related injuries below this temperature. By contrast, freeze avoidant insects cannot survive at temperatures below their SCP. Thus, the relationship between the SCP and injury/mortality is critical to determining the cold tolerance of these insects (Sinclair et al., 2015).

Cold tolerance traits, like the CT_min_, SCP, or survival following a cold stress frequently correlate strongly with insect distribution (Bozinovic et al. 2011; Gouveia et al. 2014; Andersen et al. 2015). For example, the CT_min_ and CT_max_ of *Drosophila* species have been used to accurately predict their current distributions(Overgaard et al., 2014). Similarly, the SCPs and potential for mortality at more mild temperatures were used to understand and predict the potential invasion limits of pine beetles in North America (Ungerer et al., 1999; Bleiker et al., 2019). However, given the limited number of taxa in which these relationships have been explored, we have little to no ability to generalize predictions about how climate influences species’ distributions via low temperatures to other taxa (Ouimette 2018).

Species distribution modelling is a commonly used approach to evaluate recent range shifts and forecast future shifts due to climate change. For example, they have been used to predict invasive species’ distributions (Jiménez-Valverde et al., 2011), define species’ ranges and predict impacts of climate change on available habitat (Araujo et al. 2005). Using correlations between georeferenced occurrence records and a set of environmental variables with geospatial data, these models predict a species’ suitable habitat (Elith et al., 2011). While the approach is useful for many applications, these models often violate key assumptions, such as predicting habitat suitability in novel conditions and omitting key biotic variables known to influence geographic distributions such as species interactions (Tingley et al. 2014; Briscoe et al. 2019).

One way that has been demonstrated to improve the accuracy of these models is to incorporate physiological variables or stress tolerance thresholds (e.g., heat tolerance, metabolic needs, thermal limits; (Kearney & Porter, 2009; Overgaard et al. 2014). By using variables derived directly from these physiological traits, the underlying processes explaining the species’ distribution are thought to be better incorporated into the model (Kearney & Porter, 2009). For example, Elith et al. (2010) built a distribution model for the cane toad (*Bufo marinus*) based on the impact of topography on the above-ground activity of adults and the temperature requirement of larval growth and survival. Models produced in such a way are called mechanistic or process-based models. They calculate the likelihood of presence by eliminating regions where the conditions are detrimental to species persistence (Buckley et al., 2010). In some contexts, these models have been shown to be more accurate than (Peterson, 2011), or complement weaknesses of (Martinez 2015), correlative models when modelling the fundamental niche and can strengthen predictions about future distributions (Buckley et al., 2011; Martinez 2015; Evans et al. 2016; Kotta et al. 2019). The main limitation with this approach is that experimentally derived variables can be time-consuming to develop and thus not feasible to do for large assemblages of species or over broad ranges (Peterson 2011; Evans et al. 2016). It can also be difficult to find traits that are predictive and functional in the species’ environment (Kearney & Porter, 2009).

Here, we use a widespread butterfly, the Giant Swallowtail, *Papilio cresphontes*, as a case study to test the hypothesis that low temperatures during autumn are the limiting factor of the northern edges of insect species’ distributions. Specifically, the two main objectives are to: i) find relevant cold tolerance thresholds of *P. cresphontes* collected in autumn; and ii) determine the relative importance of these thresholds on the distribution of *P. cresphontes* at the landscape level. Simultaneously, we aim to determine the cold tolerance strategy of larvae at the northern range limit. Determining the species’ cold tolerance strategy improves our knowledge about the importance of low temperatures on *P. cresphontes* survival and helps identify which low temperature thresholds are likely to be most relevant at the species’ northern range. In this study, we focus on the pre-overwinter life stage (i.e., larval stage), which is in contrast to most other studies which have determined the cold tolerance strategy of the life stage that overwinters (Radchuk et al., 2013). To determine the importance of low temperatures at a broad scale, we model the geographic distribution of *P. cresphontes* with the cold tolerance parameters we generated experimentally.

*P. cresphontes* has undergone a rapid expansion over the past decade and now occurs as far north as Ottawa, Ontario, Canada. Finkbeiner (2011) hypothesized that the range expansion into New York state from 2000 to 2010 was due to the disappearance of frost (as defined meteorologically) in September. In spite of this, they still showed frost resistance in larvae and that individuals could survive exposure to −3.9°C. However, these observations were based on field observations of only a few specimens and the cold tolerance strategy was not determined. Therefore, a more in-depth study of the impact of low temperature on *P. cresphontes* is warranted.

To establish the larval cold tolerance strategy, we measured three common cold-induced physiological thresholds: SCP, CT_min_, and survival following cold exposure. To gain a better estimate of the cold tolerance strategy, thresholds (SCP and low temperature survival) were measured for two generations and multiple sites across a latitudinal gradient at the northern range limit. Latitude has the potential to affect cold tolerance due to its correlation with climate (e.g., photoperiod, temperature; Sømme, 1982; Tanaka, 1996) but see Yoshio & Ishii, 2001). Therefore, latitude could affect when and to what extent organisms are cold hardy. Combined, these experiments were used to identify a potentially relevant low temperature threshold of larvae collected at the northern range edge. We examined the importance of low temperatures at a broad scale by comparing species distribution models of *P. cresphontes* based only on environmental data derived from other sources to models that also included the cold tolerance parameters generated experimentally.

## Materials and methods

### 1. Study system

The Giant Swallowtail (*Papilio cresphontes*) butterfly is a member of the Papilionidae family. The species’ range extends from Costa Rica throughout the North American continent. The bulk of the population is concentrated in the Eastern United States. It ranges as far north as the Ottawa region, Ontario, Canada. (Figure 1)

**Figure 1.**
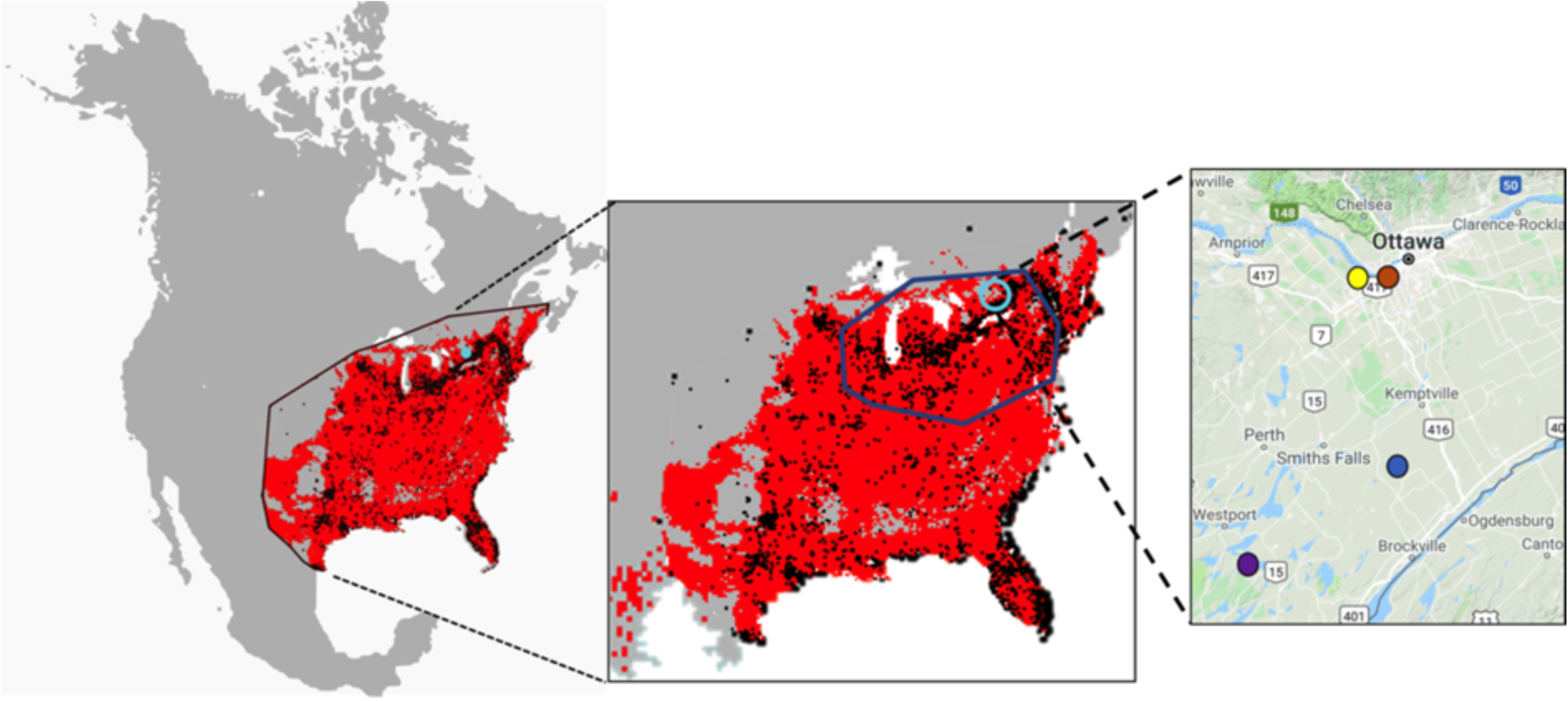
The geographic distribution and northern range of *Papilio cresphontes*. The first panel shows the entire range of *P. cresphontes* encompassed by the minimum convex polygon (thin black line). The second panel shows the position of the northern range, denoted by the solid blue line, relative to the entire range. The blue circle indicates the locations of the field sites. In the first two panels, the occurrence points (black) are shown against the predicted habitat suitability (red). The third panel shows the field sites where larvae were collected at the northern range limit: the blue dot indicates the Brockville site (44.84952, −75.75226), red indicates the Queen’s University Biological Station site (44.56747, −76.32454), brown indicates the Mud lake site (5.37192, −75.79451) and yellow indicates the Shirley’s Bay site (45.36546, −75.88302).

*P. cresphontes* larvae undergo five instars before entering the chrysalis stage (Bullock, 1991). In Ontario, there are two generations with flights occurring from May to July and again from late July to late September (Layberry, Hall & Lafontaine, 1998). The larval stage lasts from 3-4 weeks and pupae formed in the summer emerge after 10-12 days, but those that develop in autumn will remain as pupae until spring. The overwintering pupae undergoes winter diapause, a state of lowered metabolism, with reduced respiration, and no feeding or growth occurs (Scott, 1997). Once eclosed, adults live for 6-14 days (Layberry, Hall & Lafontaine, 1998).

*P. cresphontes* is known to use plants of the Rutaceae family as their primary food source in the larval stage. In Ontario, those plants consist mainly of Northern Prickly Ash (*Zanthoxylum americanum*) and Hoptree (*Ptelea trifoliata*). The adults are generalists and will gather nectar from most flowering plants, for example, goldenrod (*Solidago*), swamp milkweed (*Asclepias incarnata*) and azalea (*Rhododendron azaleastrum*)(McAuslane, 2004).

### 2. Cold tolerance experiments

#### i) Experimental overview

To determine the impact of low temperatures on *P. cresphontes* larval survival and developmental success, three experiments were conducted (Figure 2; Figure S1). The first experiment was done to determine the cold tolerance strategy of the larvae by measuring the SCP in relation to cold survival. Additionally, since the results from this initial experiment were not enough to clearly define the cold tolerance strategy, further low-temperature survival assays (i.e. the ambient exposure of larvae to low temperatures for an extended period of time) were conducted to better determine the impacts of prolonged low temperature exposure on survival and developmental success. A third experiment was conducted to measure the CT_min_.

**Figure 2.**
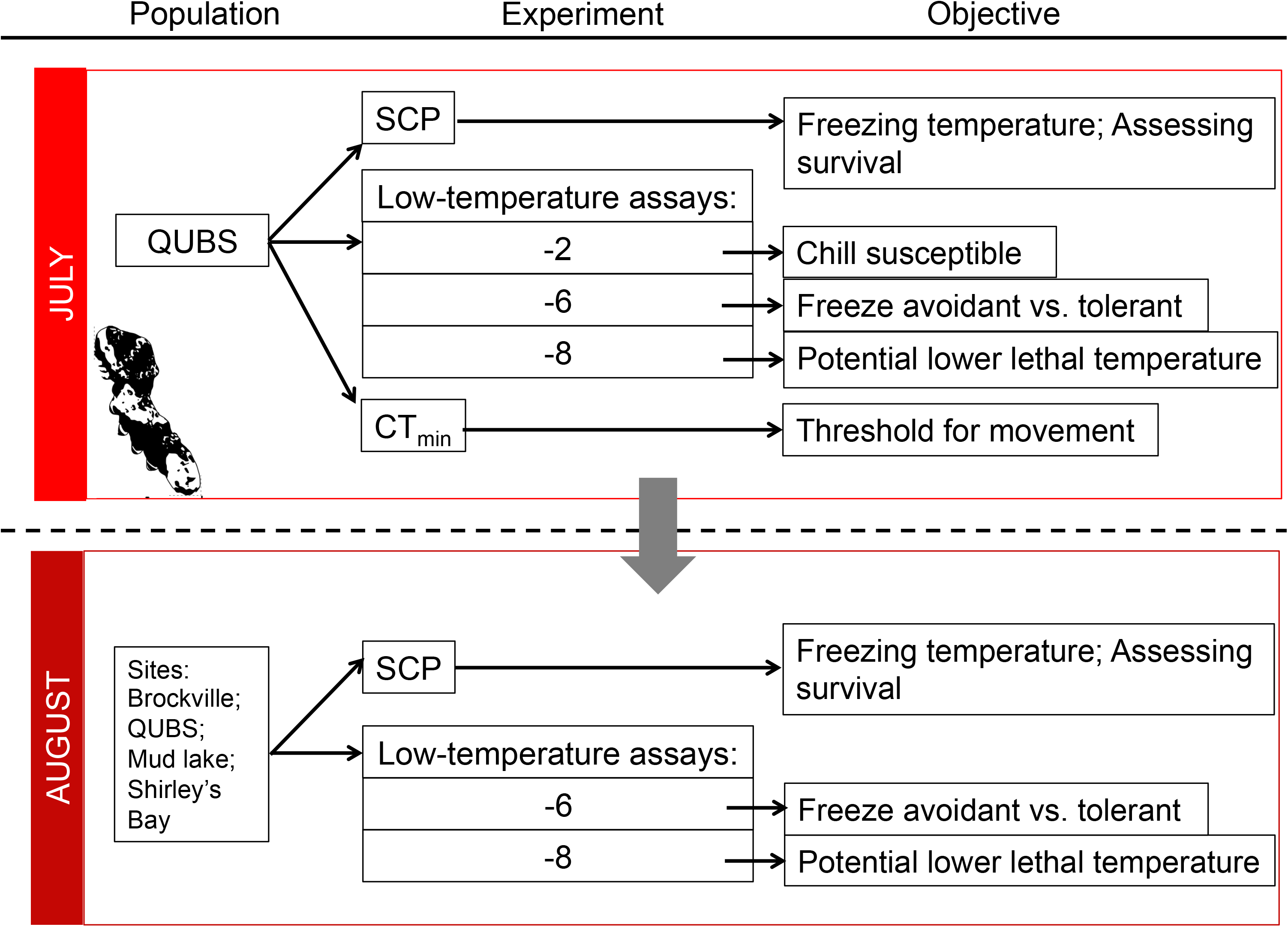
Overview of cold tolerance experiments conducted in this study. Shown are the supercooling point (SCP), low temperature survival assays, and critical thermal minimum experiments (CT_min_). Sites included in the experiments are: Brockville, Queen’s University Biological Station (QUBS), Mud lake and Shirley’s Bay. For number of larvae in each experiment, please refer to Table S1.

The SCP and low-temperature survival assays were measured for both generations and multiple sites (n=4) across a latitudinal gradient at the northern range limit (Figure 1, 2).

#### (ii) Field sampling

To gain a better understanding of the cold tolerance strategy, larval sampling occurred during the two *P. cresphontes* generations: July and August of 2018. A total of 117 larvae were collected from four sites around Ottawa, Ontario, Canada (Mud Lake, Shirley’s Bay, Brockville and Queens University Biological Station (QUBS)) spanning a 100 km latitudinal gradient at the northern range limit of *P. cresphontes* (Figure 1). These four sites were accessible and had a high abundance of larvae.

Until the start of the experiments, captured individuals in July were provided with a fresh supply of *Z. americanum* and kept in an LTCP-19 Biochamber at the University of Ottawa, Canada (45.423325, −75.683177) on a 21°C/25°C 15h: 9h L:D cycle with light intensity peaking at noon. These conditions were chosen to match the average environmental conditions in July for the Ottawa region. To replicate the conditions required for larvae to prepare to overwinter, the chamber parameters were modified weekly for the August generation to match the conditions from August to October (meteomedia.ca; Figure S1). Over the time spent in the chamber, peak daily temperatures reached a maximum 25 °C and a minimum of 15°C overnight at the beginning and by the end, daytime temperatures only reached 10°C and a minimum of 6°C overnight (Figure S1). Likewise, photoperiod fell from 14.5h to 11h. To validate our treatments, we recorded the realized climate regime in the environmental chamber using a temperature logger (HOBO Pendant, Onset Computer Corporation, Bourne, MA).

#### (iii) Experimental details

##### Experiment 1: Supercooling point (SCP) test

To identify the cold tolerance strategy of *P. cresphontes* larvae, we quantified the SCP and survival following a freezing event in the first generation (i.e. the July generation; n=27 from QUBS; Figure 2; Figure S1). The SCP test was done following the recommended methodology of Sinclair (2015) and details can be found in Appendix S1.

To get a more accurate estimate of SCP, another SCP test was done on the second generation (i.e. the August generation; n=29; Figure 2, Table S1; Figure S1). The individuals from the August generation were subjected to the same cooling protocol as described above but were kept submerged until all of the larvae had frozen. This was done to obtain the full distribution of SCP values.

##### Experiment 2: Low-temperature survival assays

To identify the consequences of chronic low temperature likely to be experienced by the larvae before pupation, three low-temperature survival assays were conducted (Figure 2; Figure S1). The duration and temperature of the assays were chosen based on the SCP determined in the first experiment and on meteorological data in the Ottawa area representing the developmental period before pupation. In the Ottawa region, this period corresponds to the month of October. We considered temperatures from 2012 to 2017, which correspond with the timing of the recent range expansion of *P. cresphontes* into the Ottawa area (Ontario Butterfly Atlas: www.ontarioinsects.org/atlas_online.htm) and used the ‘OTTAWA CDA RCS’ weather station (45.38, −75.71). For this timeframe, the mean and lowest temperatures recorded during frost events (as determined meteorologically: the span of time in which the temperature is below 0°C for more than one hour) were −1.33°C and −6.3°C, respectively, and the average frost time was 6.24 hours. Consequently, and in conjunction with the measured SCP of −5.6°C determined in the first experiment, trials were run at −2°C, −6°C, −8°C for 7 hours each.

To determine whether larvae were chill susceptible, the first assay was done at −2°C (Figure 2). Mortality at this temperature (i.e., above the SCP; see results) would indicate this species is chill susceptible whereas survival would confirm the ability of larvae to tolerate exposure to low temperatures above their SCP, thus making them chill tolerant or freeze avoidant. The second test was done at −6°C to distinguish between freeze avoidant and freeze tolerant strategies (Figure 2). Considerable mortality near the mean SCP would mean larvae are freeze avoidant whereas high survival would mean they are more likely freeze tolerant. Finally, to find a potentially ecologically relevant low temperature threshold causing high mortality (i.e. a potential lower lethal limit), we also tested larval survival at the lowest recorded temperature for October at this weather station from 2012-2017 (i.e. −8°C). Exposure at this temperature tests the maximum larval cold tolerance and their resistance to freezing since it is well below the mean SCP obtained in the previous experiment (i.e., −5.77 °C; Figure 2).

The low-temperature assays were done in an environmental growth chamber (Biochambers model LTCB-19) for both larval generations and across multiple sites (Figure 2; Table S1). The −2°C test was not repeated in August because i) there was high survival with the July individuals (see results); ii) a low number of specimens were collected in the field, and iii) this temperature was unlikely to be a limiting factor for larval survival and thus was not a priority to test. Larvae were weighed and measured before being placed in individual containers inside the chamber. The chamber temperature was brought down from 21°C to the predetermined test temperature at a rate of 0.1°C per minute. Once the desired minimum temperature was reached, the temperature remained constant for seven hours before being increased back to 21°C at the same rate. Larvae were then left to recuperate. They were allowed to feed on freshly harvested *Z. americanum* leaves during the trial and were checked daily for mortality. Death was determined when larvae did not react to physical stimuli: larvae were poked with a stick, sprayed with water and then shaken. All larvae that survived to pupation were kept until emergence.

##### Experiment 3: Critical thermal minimum (CT_min_)

To identify the temperature at which larvae lose coordination, a CT_min_ test was done on twenty larvae were caught in July from the QUBS site, following the method of Andersen et al (2014)(see Appendix S1 for details). The CT_min_ test was only done on July individuals since it required a higher number of specimens than the other tests and our sample size was limited.

#### iv) Statistical analysis

To compare survival rates of larvae of the July generation that froze to those that did not freeze during the SCP experiment, a χ2 test was used. The rate above and below the SCP for the August generation could not be compared since all larvae were exposed to low temperatures until SCP was reached (i.e. no survival rate was calculated above SCP). To test whether mean SCP differed across sites for the August generation, an ANOVA was run. Since there was no significant difference in mean SCP across sites (df=3, F-value=0.94 p = 0.44), site-level data were pooled. The assumptions of normality and homogeneity were evaluated using the Shapiro-Wilk test and the Brown-Forsythe for homogeneity of variance test, respectively.

To compare the larval survival rate and the rates of pupation and adult emergence between generations and sites for the low-temperature assays, a χ2 test was used. There was no impact of site on survival rate or developmental success in any assay (Table S2), so site-level data were pooled. All analyses were conducted in R (version 3.4.1).

### 2. Distribution modelling

#### i) Overall approach

To determine the limiting factors of the current geographic distribution of *P. cresphontes*, we modelled its range at two different spatial extents: northern range and the entire distribution (Figure 1); and using two different modelling approaches. We wanted to determine whether the relative importance of variables differs between the northern range and the entire distribution. We defined the northern range as the area encompassing the upper 50^th^ percentile of occurrences based on both latitude and longitude in the northeastern part of the range (Figure 1). The 50^th^ percentile represented a clear threshold in the latitudinal distribution of occurrences. To assess if the physiological-based cold tolerance metrics were significant predictors of *P. cresphontes’* distribution, the modelling was done using two approaches: correlative and mechanistic. The correlative approach was based on environmental data derived from other sources (e.g., remote sensing, weather stations) whereas the mechanistic approach additionally included parameters derived from the cold tolerance experiments.

For both approaches, we tested the importance of the cold tolerance strategy parameters by comparing i) model fit with and without them and ii) the proportion of suitability explained among a number of climate-related factors hypothesized to constrain the range of *P. cresphontes* at the landscape scale.

#### ii) Data

Models were built using all available occurrence records for North America from e-butterfly (www.e-butterfly.org, accessed July 2018, 2255 records), the Global Biodiversity Information Facility (GBIF; https://www.gbif.org, accessed September 2018, 3420 records) and Moths and Butterflies of North America (www.butterfliesandmoth.org, accessed October 2018, 1206 records) from 1980 to 2018. This timeframe was chosen to match the timeframe of the available environmental variables. Records that were obviously inaccurate (e.g., in the middle of the ocean) or had no geographical coordinates were removed. To reduce the likelihood of an occurrence being based on the same individual being observed on multiple occasions, duplicate records across data sources were also removed. To reduce sampling bias, we used a low-end estimate of dispersal ability (1 km) and randomly selected a single occurrence across data sources per 1 km^2^. In total, 3510 and 1457 occurrences were used for the North American and the northern range extent, respectively. Occurrences were then mapped onto a North America Lambert Conical Conic projection.

Sixteen environmental variables were initially considered for the model building based on previous work done on butterflies (Appendix S1; Table S3). The variables included represented both the direct and indirect effects (i.e. via a species’ interaction) climate can have on a species’ range (Appendix S1). Variables were created for both range extents and then screened for collinearity (Appendix S1; Table S4). All climatic data were downloaded at a 1 km resolution and restricted to 1980-2010. All data was downloaded at a North American extent, with the exception of Normalized Difference Vegetation Index (NDVI), which was global in extent.

For the mechanistic models, we created variables for both range extents based on the results from the cold tolerance experiments and minimum daily temperature data from Daymet (Table S3). Three variables were built for each extent by summing the number of days below the following thresholds: 2.14°C (i.e., CT_min_), −6.6°C (i.e. the SCP from the August generation) and −8°C (i.e. potential lower lethal limit) for each year of the study period 1980-2010 (Table S3). The SCP from the August generation was chosen since it is the one most likely to match when larvae are more likely to experience low temperatures.

#### iii) Modelling

Models were built using Maxent (version 3.4.1, Philipps et al. 2006) with the BIOMOD2 package (Damien Georges) in R (version 3.41.1). Maxent is a commonly used species distribution modelling technique for presence-only data (Merow et al., 2013). Maxent uses a maximum-entropy approach to model species distributions. It relates the species’ presences and pseudo-absences to the environmental variables to predict where the habitat is suitable across the mapped geographical space. Maxent is one of the more common species distribution modelling software (Elith et al., 2011) and it performs very well compared to other modelling techniques when dealing with presence-only data (Hernandez et al., 2006). Details on the modelling are presented in Appendix S1.

#### iv) Statistical analysis

To determine the role of low temperature in limiting the distribution of *P. cresphontes*, model accuracy was compared across the two extents and approaches using t-tests. Comparisons were based on AUC, Kappa and TSS.

## Results

### 1. Cold tolerance experiments

#### Experiment 1: Supercooling point (SCP) test

The results from the July and August SCP experiments provide support for a chill susceptible strategy in *P. cresphontes*. The mean SCP for July was −5.8°C (0.2SE; n=14). There was no significant difference in survival for July larvae that froze compared those that did not freeze during the SCP experiment (above: 77% (10/13) vs. below: 43% (6/14 larvae); χ2= 1.2, df = 1, p-value = 0.2).

The mean SCP for all August larvae was −6.6°C (0.2SE; n=28) and for the upper 50% of larvae (directly comparable to the July experiment), the mean SCP was −5.9°C (0.2SE; n=14). The August larvae had a 3.5% (1/29) survival rate following freezing (i.e., exposure to temperatures below the mean SCP). The low survival below SCP and apparent lack of change in the SCP suggests that larvae are tolerant of moderate chilling, but intolerant of freezing, and are not undergoing a change in SCP under simulated autumn conditions.

#### Experiment 2: Low-temperature survival assays

##### i) Survival

Our low-temperature survival assays further demonstrate that larvae are chill tolerant or modestly freeze avoidant, as they are unable to survive exposure to temperatures below their SCP. In the −2°C test, July larvae had a survival rate of 93% after 24 hours (Table 1). Therefore, single ecologically-relevant exposures to temperatures above their SCP do not have a large impact on larval survival. Moreover, while the survival rate of larvae in July was more affected by exposure to −6°C than −2°C (Table 1), it was still rather high (70%). Therefore, larvae are able to survive a 7 h cold exposure near their SCP and thus may be chill tolerant or freeze avoidant.

**Table 1.** Survival rates and developmental success for the low-temperature assays. Test temperature and generation are shown. The percentage of initial larvae that survived the first 24 hours after the experiment, successfully pupated and eclosed as adults is alos shown. All percentages are based on the number of larva that began the experiment.

| Test | Generation | Number of larvae | Larval survival (%) | Pupation (%) | Eclosion (%) |
| --- | --- | --- | --- | --- | --- |
| -2°C | July | 23 | 93 | 78.5 | 52 |
| -6°C | July | 10 | 70 | 60 | 30 |
|  | August | 17 | 100 | 100 | 0 |
| -8°C | July | 8 | 12.5 | 12.5 | 12.5 |
|  | August | 10 | 10 | 10 | 0 |

**Table 2.** Contribution of the environmental variables in explaining habitat suitability across different model extents and approaches. Shown is the rank order of variable importance and mean (± S.E) proportion of variance explained. In bold is the variable that explains the most amount of variation for each model type.

| Variables | North America |  |  |  | Northern range |  |  |  |
| --- | --- | --- | --- | --- | --- | --- | --- | --- |
|  | Correlative |  | Mechanistic |  | Correlative |  | Mechanistic |  |
|  | Rank order | Variation explained (SE) | Rank order | Variation explained (SE) | Rank order | Variation explained (SE) | Rank order | Variation explained (SE) |
| **Growing degree-days** | **1** | **31.33 (0.19)** | **1** | **27.34 (0.23)** | **1** | **37.17 (0.08)** | **1** | **35.03 (0.06)** |
| Precipitation | 2 | 24.87 (0.08) | 2 | 21.70 (0.085) | 2 | 23.94 (0.08) | 2 | 22.47 (0.07) |
| Extreme maximum temperature | 3 | 17.54 (0.06) | 3 | 16.22 (0.06) | 6 | 2.38 (0.02) | 7 | 2.41 (0.02) |
| Normalized Difference Vegetation Index | 4 | 15.45 (0.10) | 4 | 15.87 (0.11) | 4 | 16.77 (0.11) | 4 | 14.89 (0.11) |
| Mean temperature of the coldest month | NA | NA | NA | NA | 3 | 16.9 (0.06) | 3 | 15.57 (0.05) |
| Precipitation as snow | 5 | 10.80 (0.07) | 5 | 10.05 (0.07) | 5 | 2.84 (0.02) | 6 | 3.46 (0.02) |
| CT<sub>min</sub>* | NA | NA | NA | NA | NA | NA | 5 | 5.97 (0.03) |
| Potential lower lethal temperature* | NA | NA | 6 | 8.80 (0.04) | NA | NA | 8 | 0.16 (0.004) |

\* variable derived experimentally

August larval survival was significantly higher than that of the July larvae following exposure to −6°C (100%, 17/17; χ^2^ −squared = 5.1, df = 1, p-value = 0.024; Table 1). This finding provides further evidence in support of a modestly freeze avoidant strategy since the mean SCP for the August generation (−6.6°C) was lower than the test temperature (−6°C) and survival was still quite high.

In comparison, in the extreme low-temperature test (−8°C), larvae had marginal survival in both generations (July: 12.5% (1/8); August: 10% (1/10)) and there was no improvement in survival across generations (χ^2^ = 3.03e-31, df = 1, p-value = 0.99; Table 1). This low survival further suggests larvae may use a freeze avoidant strategy since this temperature is below the mean SCP measured in the August generation.

##### ii) Developmental success

Overall, low temperature exposure impacted pupation and adult eclosion. The percentage of larvae that successfully pupated and eclosed generally decreased as the temperatures got lower (Table 1). The percentage of larvae that successfully pupated at −8°C was low (July: 12.5%; August: 10%). Therefore, although larval survival can be high after exposure to low temperatures, chilling also had effect on developmental success, which is indicative of sub-lethal chilling injury occurring, even in the absence of freezing.

#### Experiment 3: CT_min_

The average CT_min_ of larvae was 2.14°C (0.26SE; n=20).

### 2. Distribution modelling

#### i) Model performance

Model predictive ability varied depending on the metric of accuracy used. All models were considered good based on TSS (i.e. between 0.4-0.75) and AUC (>0.8), whereas based on Kappa, the northern range models were weak (<0.4) and the full range models were fair (>0.5; Figure 3).

**Figure 3.**
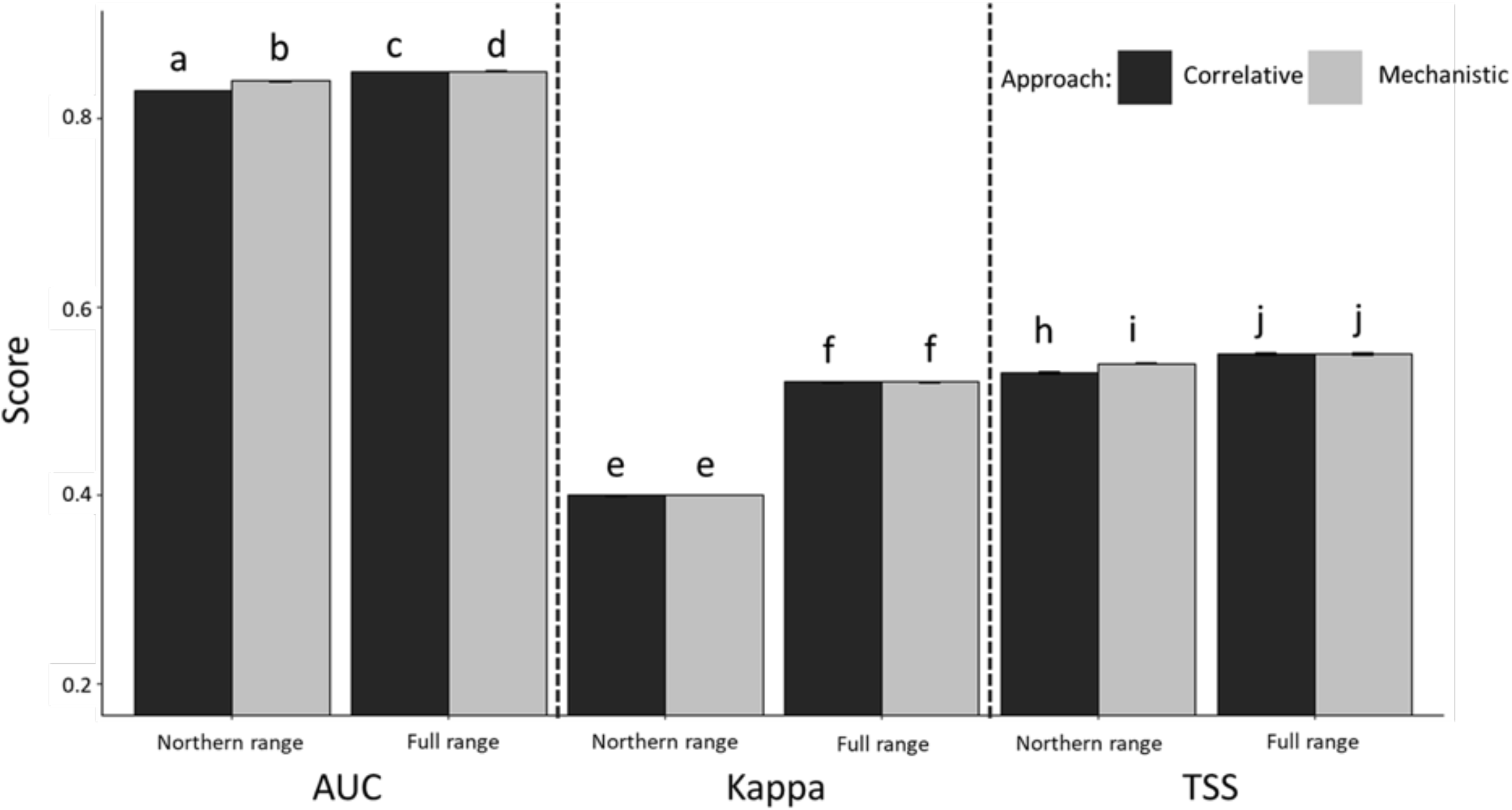
A comparison of model accuracy for the two model extents (northern and full range) and approaches (correlative and mechanistic) using all three evaluation metrics (AUC (Area under the receiver operating characteristic curve), Kappa, TSS (True skills test). Mean accuracy scores and standard error bars across 100 iterations are shown with the y-axis cropped at 0.2. Grey and black bars represent the mechanistic and correlative models, respectively. Letters indicate that mean scores are significantly different from each other (p<0.05).

Statistically, the mechanistic models were significantly more accurate than the correlative models in some but not all cases (Figure 3; Table S5). However, in the select cases where there was a significant difference between modeling approaches, the effect size was small, implying the difference is unlikely to be biologically significant (e.g., AUC difference for northern range comparison: 0.011; Figure 3, Table S5). Therefore, it is unlikely the inclusion of cold tolerance metrics significantly improves the prediction of habitat suitability of *P. cresphontes*.

The accuracy of the full range models was significantly higher than the accuracy of the northern range models for all evaluation metrics and approaches (Figure 3; Table S6). Although the effect size is also not large, the significant difference in accuracy was consistent across all comparisons (Figure 3; Table S6). Therefore, the habitat suitability of *P. cresphontes* is predicted more accurately across its full range than its northern range.

#### ii) Variable importance

Growing degree-days was the most important factor explaining the habitat suitability of *P. cresphontes* across all model extents and approaches, explaining between 27-35% of the variation in habitat suitability (Table 3). Precipitation was the second most important factor across both extents and approaches. Across the full and northern range extents, GDD and precipitation combined explained the majority of the variation in habitat suitability (~50%; Table 3). The physiologically derived variables included in the final model (i.e., CT_min_ and the potential lower lethal limit) only explained a small part of variation in habitat suitability (~7.5%) and ranked 5^th^ and 6^th^ in model contribution (Table 3).

## DISCUSSION

Although insects are likely to be particularly vulnerable to low temperatures during autumn, no study, to our knowledge, has considered low temperatures during autumn as a limiting factor on the geographic distributions of insects. Here, we test this hypothesis at the northern edge of the distribution of the widespread butterfly, the Giant Swallowtail, *P. cresphontes*. Our study contributes three main findings that lead to a rejection of this hypothesis. First, given the survival of larvae to prolonged exposures to temperatures close to but not below their SCP, they are chill tolerant or modestly freeze avoidant at their northern range limit at this time of year. Their low survival below the SCP provides support that this species is unable to handle freezing before pupation and that they may use a freeze avoidant strategy. However, further testing for seasonal plasticity in the SCP is required to confirm a freeze avoidant strategy as we had limited power and were unable to directly compare the change in SCP between the two generations. An active depression of the SCP over the season would indicate that larvae are accumulating cryoprotectants and that the larval cold tolerance could be increasing, thus supporting a freeze avoidant strategy.

Our results demonstrate that ecologically relevant exposures to temperatures above the SCP are not lethal, but below the SCP are; thus providing a clear picture of the lower end of thermal tolerance for this species. This means that for *P. cresphontes* larvae, a single overnight exposure to temperatures near −8°C will impede their survival and could represent a potential lower lethal limit. Consequently, areas with early autumn temperatures that reach −8°C should be inhospitable for *P. cresphontes*. However, in the Ottawa region, which is at the northern range limit, there are only 0-3 frost episodes on average from September to October. Therefore, low temperatures (i.e. <-7°C) during early autumn are unlikely to be experienced by the larvae there or anywhere in its current range. This suggests that a single exposure to low temperatures in early autumn is unlikely to limit the current northern range of *P. cresphontes* through cold-induced mortality. However, there may be sublethal effects of chilling (e.g., effects on developmental success), which could still impact overall insect fitness.

Second, we found that exposure to normal frost temperatures for this time of year in this area (i.e., −1°C to −2°C), which are above the SCP, did not affect larval survival. Therefore, it is also unlikely that frost in early autumn is a limiting factor of *P. cresphontes’* northern range. This is in concordance with the field observations from Finkbeiner (2011), which showed that *P. cresphontes* larvae can survive frost events. Since larvae had no problem reaching pupation after exposure to −2°C, the most common frost temperature, and the majority of larvae still successfully pupated (60%) after the low frost temperature test (i.e., −6°C), the cold tolerance of larvae at the current northern range limit seems sufficient to cope with the climate they experience. However, further testing in ambient conditions and with a proper control is needed to determine whether cold exposure on larvae have significant sub-lethal effects on eclosion as rates of eclosion in this experiment were less than 30% but a control group was not included due to sample size constraints. It has been demonstrated elsewhere that the success of each life stage matters in the response of butterfly species to climate change (Radchuk et al. 2012).

Third, the species distribution modelling results were consistent with the experimental results: cold related variables did not explain the distribution of *P. cresphontes* at a broad scale. Together, these results provide strong support that exposure to temperatures above −6.6°C during autumn does not limit the northern range of *P. cresphontes*. Instead, growing degree-days and precipitation are the most important predictors, of those tested here, on the distribution of *P. cresphontes* at a broad scale. While other factors were not tested here, for example, biotic interactions (e.g., host plant occurrence), our results suggest that *P. cresphontes* depends on specific heat accumulation and water availability to complete its life cycle. These factors have also been identified as important in predicting the range of other butterflies (Luoto et al., 2006; Eskildsen et al., 2013). Evidence suggests there is a strong relationship between the number of growing degree-days and the growth rate of larvae, and the foraging activities of adults (Ritland & Scriber, 1985; Kukal & Dawson, 1989; Schneider & Root, 2002). Precipitation can limit the range of insects directly due to dehydration or indirectly by imposing limitations on primary productivity, which in return transposes to higher resource availability.

The full range model was more accurate at predicting the distribution of *P. cresphontes* than the northern range model. Similar results have been found for other species, with model accuracy and performance increasing with larger study extents (VanDerWal et al., 2009; Connor et al., 2019). This is likely a result of smaller environmental gradients in the northern range, which causes the model to have a greater difficulty detecting differences between presences and absences, and in turn leads to difficulties predicting presences and absences in novel areas (Smith & Santos, 2019). Prevalence can also decrease with larger study extents (Barbet-Massin et al., 2012). However, the accuracy of the two models was still different using TSS, a metric that is not sensitive to prevalence (Table S4; Figure 3). Nevertheless, even if model performance was lower in the northern range relative to the full range, the model was still valid and performed adequately across all metrics.

While growing degree-days and precipitation explain a lot of the variation in habitat suitability for *P. cresphontes*, we may have underestimated the role of cold tolerance at the northern range edge for three reasons. First, low temperature thresholds, like the CT_min_, although not lethal, could still impact *P. cresphontes’* survival in natural conditions due to the enhanced risk of predation or starvation linked with the loss in mobility at temperatures below 2°C. While a possibility, this hypothesis depends on a high likelihood of predation, which is unknown across *P. cresphontes’* range (Hazel et al., 1998; McAuslane, 2009). Low temperature may also lead to sublethal effects on developmental success, energy status and behaviour. Second, it is also possible that more prolonged or repeated exposure to low temperatures could impact *P. cresphontes* larval survival or have sublethal effects at the northern range edge because we did not test for the effects of repeated cold injury or chilling. In *Drosophila*, multiple cold events could cause an accumulation of cold injuries resulting in larval death or in sub-lethal effects such as decreased feeding, fecundity and dispersal ability (Marshall & Sinclair, 2012). Third, the SCP estimated here is effectively a theoretical one. Since we could not collect and test the last emerging larvae of the season due to the difficulty in finding them, they might not have reached their peak cold tolerance and therefore *in situ*, SCP could be lower (i.e., their cold hardiness may have been underestimated). On the other hand, the *in situ* SCP may be higher since environmental factors in natural conditions, such as air moisture or exposure to ice nucleators, may further affect survival at temperatures above SCP. Nevertheless, because the low temperature thresholds derived experimentally in this study could be an underestimate, the mechanistic models may not have been optimized perfectly, thus underestimating the role of cold tolerance metrics at a broad scale. However, since temperatures below the SCP (i.e. <-7°C) are very unlikely to occur multiple times before pupation, if at all, low temperatures during autumn remain unlikely to limit the northern range of *P. cresphontes*.

Given our lack of knowledge about the overwintering ecology and physiology of this species in its northern range, pupal overwinter survival may be a more important limiting factor for the species’ northern range limit. Unfortunately, it is unknown whether they overwinter in the leaf litter on the ground or tree branches above the ground. If on the ground, they would likely be covered in snow (average annual snowfall in Ottawa is 224 cm (Environment and Climate Change Canada)) and therefore buffered against low temperatures. The temperature 30 cm below snow cover is usually −4°C (Flin 2008). This behaviour would increase their chances of surviving the winter. Indeed, West (1996) showed that in Virginia, *P. cresphontes* pupate 3-25 cm off the ground on dead branches. In the northern range, it is likely that even at these heights, chrysalids would be covered by snow. While ontogenetic variation in thermal tolerance is present in other species (Terblanche et al. 2006; Marais et al. 2009; MacLean et al., 2016; but see Ouimette, 2018) and only larvae were tested in this study, *P. cresphontes* larvae purge their food and liquids before pupation, thus they are likely to be even more cold tolerant at the pupal stage. Other members of *Papilio* are more cold tolerant than what we determined for the larval stage of *P. cresphontes. P.canadensis* and *P.glaucus*, are considered to be freeze tolerant at the pupal stage and have an SCP of −23°C to −27°C (Kukal et al., 1991). *P. xuthus* is freeze avoidant with an SCP of −25°C (Shimada 1988). Therefore, the pupae of *P. cresphontes* are likely to have even greater cold hardiness than the larvae and thus be able to survive the winter at the northern range if they pupate at similar heights as the populations in Virginia.

### Conclusions

Our results demonstrate that the cold tolerance of *P. cresphontes* larvae is well matched to its environment in the northern part of its range and that it is unlikely that low temperatures in autumn are limiting its range. This suggests that the warming of autumn temperatures over the past decade is unlikely to have been the factor that led to the recent range expansion of *P.cresphones*, as originally hypothesized by Finkbeiner (2011). Instead, the warming of winter temperatures could have ameliorated overwintering survival, leading to an improvement in habitat suitability. Further study on determining the key factor(s) limiting the northern range of *P. cresphontes* should focus on defining the cold hardiness of the pupal stage, the overwintering behaviour and the ecological relevance of the physiological thresholds found in this study. Determining the ultimate factors that limit species’ distributions will be critical in accurately predicting species’ range shifts in response to future climate change.

## Acknowledgements

We thank S. Foster, S. Gilmour and S. Lalonde for help conducting the experiments, M. Larrivée and eButterfly for providing data, and R. Layberry and O. Eydt for their expertise. We are grateful to J. Kerr and C. Harris for providing helpful feedback on previous versions of this manuscript. Thanks to the Ontario Ministry of Natural Resources and Forestry for collection permits and the National Capital Commission for site permission to Mud Lake and Shirley’s Bay. HMK and HAM both acknowledge funding through NSERC Discovery Grants. Equipment used in the experiments was acquired through support from the Canadian Foundation for Innovation and Ontario Research Fund Small Infrastructure Fund (to HMK and HAM).

## Author’s Contributions

PT and HMK: conceived the ideas, designed the methodology and wrote the paper. PT collected data and performed analyses. HM: helped design the methodology and edit the paper. All authors contributed critically to the drafts and gave final approval for publication.

## Data availability

Should the manuscript be accepted, the data supporting the results will be archived in an appropriate public repository (Dryad or Figshare) and the data DOI will be included at the end of the article.

## Notes

### Competing Interest Statement

The authors have declared no competing interest.

### Summary of Updates

Modelling results revised and Supplemental files updated

