## Appendix for "Autumn larval cold tolerance does not predict the northern range limit of a widespread butterfly species"

**Appendix S1.** Additional methodological details related to cold tolerance experiments and species distribution modelling

Experimental details for cold tolerance experiments

*i) Experiment 1: Supercooling point test*

To identify the cold tolerance strategy of *P. cresphontes* larvae, we quantified the SCP and survival following a freezing event in the first generation (i.e. the July generation; n=27 from QUBS; Figure 2; Figure S1). The SCP test was done following the recommended methodology of Sinclair (2015). Before being cooled, the larvae were weighed with an ultra-micro Sartorius scale and placed in individual 15 ml centrifuge tubes. Body temperatures were measured using type K thermocouples in direct contact with the larvae and linked to a Pico-Tech TC-08 data logger. The vials were then suspended in a refrigerated circulator (21L with advanced controller, VWR) filled with a mixture of ethylene glycol and water, and cooled down at 0.1°C per minute from 21°C. The test lasted between 200 and 300 min until half the specimens had reached SCP. After the SCP test, larvae were then moved to individual containers inside the LTCP-19 Biochamber at temperature and photoperiod described above (section ii). Larvae were monitored daily until death or emergence. Individuals were considered dead if they did not react to physical stimuli: larvae were poked with a stick, sprayed with water and then shaken.

*ii) Experiment 3: Critical thermal minimum ($CT_{min}$)*

To identify the temperature at which larvae lose coordination, a $CT_{min}$ test was done following the method of Andersen et al (2014). Twenty larvae were caught in July from the QUBS site, weighed and measured before being placed in 50 ml Eppendorf vials. These vials were then mounted on an aluminum rack, which was submerged in a refrigerated circulator (21L with advanced controller, VWR) containing a mixture ethylene glycol and water. The bath was cooled from 21°C to -20°C at a rate of 0.2°C per minute. The tubes were periodically (i.e., approximately every 2 min) prodded using a plastic spoon during the cooling process to stimulate movement. The $CT_{min}$ was determined when individuals fell on their sides and were no longer able to stand. Larvae were kept in the bath until the final larva fell, at which point the whole rack was removed from the solution and allowed to warm back up. Larvae were then left to recuperate in a growth chamber (Biochambers model LTCB-19) at the average conditions in July, as described above. Larvae were checked daily for mortality.

2. Distribution modelling

*i) Additional details about data*

Sixteen environmental variables were initially considered for the model building based on previous work done on butterflies (Table S3; Araújo & Luoto, 2007; Finkbeiner et al., 2011; Roland & Matter 2016).Variables included low and high temperature requirements (i.e., mean temperature of the coldest month and extreme maximum temperature), precipitation, measures of heat accumulation (i.e., growing degree-days), vegetation (i.e., NDVI) and frost (i.e., temperatures below 0°C for at least one hour) (Sykes et al., 1974). As a measure of heat accumulation, GDD measures the length of the growing season, which determines the amount of time there are favourable conditions for growth and development in plants and insects (Chuine, 2010; Régnière et al., 2012). Growing degree-days (GDD) were modelled using a 10°C base (see Table S3 for calculation), which is a commonly used threshold among butterflies and other insects (Nufio et al. 2010; Cayton et al. 2015). Other variables are related to primary production and resource availability such as NDVI and precipitation. The impact of frost events were assessed since they were previously hypothesized to be a contributing factor for the range expansion of *P. cresphontes* (Finkbeiner et al., 2011). Specifically, we tested different frequencies of frost events: the number of frost-free days and duration of the frost-free period. The effect of the intensity of frost events (i.e., how cold it was during the frost event) could not be tested since the necessary climatic variables were not available.

To reduce collinearity among variables, a variance inflation factor (VIF) analysis ('car' package (John Fox)) with a threshold of 10 was conducted. This threshold was chosen instead of a more restrictive one (e.g., 3 or 5) as it provides a compromise between collinearity and model usefulness (O'brien, 2007). The test was repeated for each of the model extents and approaches (n=4). With the VIF test, the initial list of 16 variables was reduced to eight for the northern range and six for the full range model (Table S4). The final list of variables used for the correlative modelling approach was: extreme maximum temperature, precipitation as snow, precipitation, GDD, NDVI, and mean temperature of the coldest month (Table S4). Additionally, for the mechanistic models, $CT_{min}$ and the potential lower lethal limit temperature were also included.

*ii) Additional details about modeling*

Models were built using Maxent (version 3.4.1, Philipps et al. 2006) with the BIOMOD2 package (Damien Georges) in R (version 3.41.1). To model the range, we included hinge features, which allow non-linear relationships, in the model and allowed the model the possibility of clamping (i.e., the default). Clamping serves to limit how much leniency there is in the range of values projected compared to what was observed. This action is important when predicting into novel geographic space (Stohlgren et al. 2011). However, since the training data and the projection extent were the same, no clamping was done. For each model extent and approach, a background extent was created using the minimum convex polygon function in R (Package adehabitatHR v0.4.16; Calenge, 2006). Pseudo-absences were randomly generated within the minimum convex polygons (n=10000; default value).

The models were calibrated using five-fold cross-validation (i.e., the observations were divided into k groups, or folds, of equal size). In cross-validation, the first fold is treated as a validation set, and the model is fit on the remaining k – 1 folds (James et al., 2013). Iterations were limited to 5000 to leave enough time for model convergence. All other parameters were left at default values.

Models were evaluated using three metrics that assess various aspects of accuracy and discrimination (i.e., the ability of the model to distinguish between suitable and non-suitable habitat): Area Under the Curve of the receiver operating characteristics (AUC), Kappa, and True Skills Statistic (TSS). AUC characterizes the model's ability to correctly predict if a presence or absence is a true presence or absence. A value below 0.5 means that the model is no better at predicting occurrences than random, whereas a value of 1 would mean the model predicts all presences/absences perfectly (Yackulic et al., 2013). Although it is the most commonly used metric for assessing model accuracy of species distribution models (Yackulic et al., 2013), it has been criticized for being unreliable when sample sizes are small and due to its reliance on the number of background points. As such, in these contexts, AUC should mainly be used to compare models built with the same variables and occurrences.

The discrimination of the models was further tested using Kappa and TSS. These two metrics use confusion matrices to compare models' abilities to predict occurrences correctly (Allouche et al. 2006). While Kappa has been shown to be more sensitive to prevalence (i.e. the proportion of locations that are occupied), TSS provides an alternate validation metric that is independent of prevalence (Allouche et al., 2006). Kappa scores between 0.4 to 0.6 indicate fair agreement, 0.6 to 0.8 indicate moderate agreement, and values greater than 0.8 indicate strong agreement. A TSS score between 0.40 and 0.75 indicates good predictive performance of the model while a TSS score above 0.75 indicates excellent performance (Allouche et al., 2006).

Models were run 100 times, and the sensitivity (i.e., the proportion of correctly predicted presences) and specificity (i.e., the proportion of correctly predicted absences) were extracted from each iteration and used to calculate mean and standard error of AUC, Kappa, and TSS (Cerasoli et al., 2017). The proportion of variation explained by the variables was also extracted from each iteration and averaged across all runs.

**Appendix S2.** Additional results.

1. Table S1. Number of larvae among generations and sites for the experiments.
2. Table S2. A comparison of larval survival rate and the rates of pupation and adult emergence across sites for the low-temperature assays.
3. Table S3. Information about the original list of variables initially considered for the modelling.
4. Table S4. Results from the variance inflation factor (VIF) analysis used in the species distribution modelling.
5. Table S5. Comparison of the accuracy of species distribution models between approaches.
6. Table S6. Comparison of the accuracy of species distribution models between spatial extents.
7. Figure S1. Experimental details for cold tolerance tests.

Table S1. Number of larvae among generations and sites for the three experiments. The sites are: Queen's University Biological Station (QUBS), Mud lake, Shirley's Bay and Brockville.

| Generation | Experiment | Temperature treatment (°C) | Site | Number of larvae |
|---|---|---|---|---|
| July | SCP | | QUBS | 27 |
| | Low temperature | -2 | QUBS | 15 |
| | | | Shirley's Bay | 2 |
| | | | Brockville | 6 |
| | | -6 | Mud lake | 10 |
| | | -8 | Mud lake | 8 |
| | $CT_{min}$ | | QUBS | 20 |
| August | SCP (total n=29) | | QUBS | 14 |
| | | | Mud lake | 5 |
| | | | Brockville | 3 |
| | | | Shirley's Bay | 7 |
| | Low temperature | -6 | QUBS | 8 |
| | | | Mud lake | 5 |
| | | | Shirley's Bay | 3 |
| | | -8 | QUBS | 2 |
| | | | Mud lake | 5 |
| | | | Shirley's Bay | 3 |

Table S2. A comparison of larval survival rate and the rates of pupation and adult eclosion across sites for the low-temperature assays (i.e., -2°C, -6°C, -8°C tests). The results from χ2 goodness-of-fit tests are shown. The NAs are in cases where all individuals for a given test were from the same site or the test was not repeated for both generations. See Table S1 for the number of larvae across sites.

| Test | Generation | Life stage | χ2 | Degrees of freedom | p value |
|---|---|---|---|---|---|
| -2°C | July | Larval | 0.40 | 2 | 0.40 |
| | | Pupal | 2.41 | 2 | 0.30 |
| | | Adult | 0.93 | 2 | 0.63 |
| | August | Larval | NA | NA | NA |
| | | Pupal | NA | NA | NA |
| | | Adult | NA | NA | NA |
| -6°C | July | Larval | NA | NA | NA |
| | | Pupal | NA | NA | NA |
| | | Adult | NA | NA | NA |
| | August | Larval | 1.53 | 2 | 0.47 |
| | | Pupal | NA | NA | NA |
| | | Adult | NA | NA | NA |
| -8°C | July | Larval | NA | NA | NA |
| | | Pupal | NA | NA | NA |
| | | Adult | NA | NA | NA |
| | August | Larval | 4.44 | 2 | 0.11 |
| | | Pupal | NA | NA | NA |
| | | Adult | NA | NA | NA |

Table S3. Original list of variables initially considered for the modelling and their acronyms. The data source is shown as well as the calculation used to produce the final rasters. All variables are averaged over the period from 1980-2010 and the original map projection of all variables was Lambert Conical Conic except for NDVI, which was sinusoidal.

| **Acronym** | **Full name (units)** | **Data source** | **Calculation** | **Importance** |
|---|---|---|---|---|
| NDVI | Normalized Difference Vegetation Index | Modis/Terra project | Averaged monthly NDVI raster's over the time frame. | Seto 2004, Pettorelli 2005 |
| bFFP | beginning of frost free period (FFP; day of year) | Databasin | The day of the year on which FFP begins | Hayes 1982, Westwood & Blair 2010 |
| eFFP | end of frost free period (day of year) | Databasin | The day of the year on which FFP ends | Hayes 1982, Westwood & Blair 2010 |
| FFP | frost free period (days) | Databasin | The number of days between the last spring frost and the first autumn frost | Hayes 1982, Westwood & Blair 2010 |
| NFFD | number of frost free days | Databasin | Number of days above 0°C. | Hayes 1982, Westwood & Blair 2010 |
| PAS | average precipitation as snow (mm) | Databasin | Accumulated snowfall averaged across the time frame | Roland & Matter 2016 |
| Precip | average precipitation as rainfall (mm) | Databasin | Accumulated rainfall averaged across the time frame | Storch et al. 2003 |
| EMT | extreme minimum temperature (°C) | Databasin | Lowest temperature recorded for every given year, averaged over the timeframe. | Crozier 2003 |
| EXT | extreme maximum temperature (°C) | Databasin | Highest temperature recorded for every given year, averaged over the timeframe. | Malcolm et al. 1987 |
| MCMT | mean temperature of the coldest month (°C) | Databasin | Average daily temperature for the coldest month | Crozier 2004; Luoto et al., 2006 |
| MWMT | mean temperature of the warmest month (°C) | Databasin | Average daily temperature for the warmest month | Crozier 2004 |
| octtre | average temperature in October (°C) | Daymet | Average daily temperature for October. | Larsen & Lee 1994 |
| GDD | average growing degree-days of base 10°C | Daymet | ((Tmax + Tmin) / 2) -10 | Luoto et al., 2006 |
| SCP | average number of days per year below -6.6°C (days) | Daymet | Based on the experimentally derived SCP; a day was counted if the average daily temperature reached -6.6°C. Days were counted and averaged across time frame | Ungerer et al., 1999 |
| $CT_{min}$ | average number of days per year below 2.14°C (days) | Daymet | Based on the experimentally derived $CT_{min}$; a day was counted if the average daily temperature reached 2.14°C. Days were counted and averaged across time frame | Andersen et al. 2015 |
| PLLT | Potential lower lethal temperature; Average number of days per year below -8°C (days) | Daymet | Based on the experimentally derived potential lower lethal temperature; a day was counted if the average daily temperature reached -8°C. Days were counted and averaged across time frame | Andersen et al. 2015 |

Table S4. Results from the variance inflation factor (VIF) analysis conducted on the 16 climatic variables used in the species distribution modelling for the two different extents. Shown here are the variables included in the final models based on a threshold of 10 (i.e., those with a score below 10). Variables that were 'excluded' had a VIF above 10.

| Extent | Variables | VIF values |
|---|---|---|
| Northern range | Normalized Difference Vegetation Index | 2.03 |
| | Extreme maximum temperature (°C) | 3.08 |
| | Precipitation as snow (mm) | 1.16 |
| | Growing degree-days | 5.55 |
| | Potential lower lethal temperature (days)* | 1.53 |
| | Mean temperature of the coldest month (°C) | 5.59 |
| | Precipitation (mm) | 1.77 |
| | $CT_{min}$ (days)§ | 6.95 |
| Full range | Normalized Difference Vegetation Index | 2.48 |
| | Extreme maximum temperature (°C) | 3.59 |
| | Precipitation as snow (mm) | 2.80 |
| | Growing degree-days | 2.02 |
| | Potential lower lethal temperature (days)* | 1.21 |
| | Precipitation (mm) | 3.31 |

*Average number of days per year below -8°C

§Average number of days per year below 2.14.

Table S5: Comparison of the mean accuracy of species distribution models with and without mechanistic variables at two spatial extents: northern range and full range. Shown are the t-test results comparing AUC (the area under the receiver operating characteristic curve) and TSS (true skill statistic). The comparisons in bold are statistically significant (p<0.05). Differences in scores are visually represented in Figure 3.

| Extent | Metric | Mean difference in score* | t-value | p-value | Degrees of freedom |
|---|---|---|---|---|---|
| Northern range | **AUC** | **0.011** | **2.59** | **0.01** | **999.32** |
| | Kappa | -0.0101 | 0.21 | 0.84 | 999.81 |
| | **TSS** | **0.077** | **4.75** | **2.4e-06** | **999.23** |
| Full range | **AUC** | **0.0027** | **18.32** | **2.2e-16** | **999.95** |
| | Kappa | 0.0012 | 2.74 | 0.06 | 999.59 |
| | TSS | 0.038 | 1.58 | 0.12 | 999.41 |

* Mean= mechanistic score - correlative sore

Table S6: Comparison of the accuracy of species distribution models between the two extents (northern range and full range) and for both approaches (correlative or mechanistic). Shown are the t-test results comparing AUC (the area under the receiver operating characteristic curve) and TSS (true skill statistic). The comparisons in bold are statistically significant (p<0.05). Differences in scores are visually represented in Figure 3.

| Approach | Metric | Mean difference in score* | t-value | p-value | Degrees of freedom |
|---|---|---|---|---|---|
| Correlative | **AUC** | **0.019** | **8.36** | **5.3e-16** | **957.11** |
| | **Kappa** | **0.095** | **109.94** | **2.2e-16** | **958.51** |
| | **TSS** | **0.023** | **6.61** | **6.6e-11** | **957.4** |
| Mechanistic | **AUC** | **0.011** | **24.33** | **2.2e-16** | **977.72** |
| | **Kappa** | **0.11** | **125.72** | **2.2e-16** | **950.57** |
| | **TSS** | **0.019** | **4.75** | **2.4e-06** | **958.14** |

* Mean= Full range score – Northern range score

Figure S1. Experimental details for cold tolerance tests. (a) Overview of starting and testing temperatures and dates of collection and tests for July generation. Only first collection date and only rough test dates are shown to improve visualization. (b) Profile of temperature conditions in the environmental chamber for larvae from the August generation. Shown is the diurnal temperature range the chamber was programmed for a given week. Parameters were modified weekly to match the conditions from August to October (meteomedia.ca).

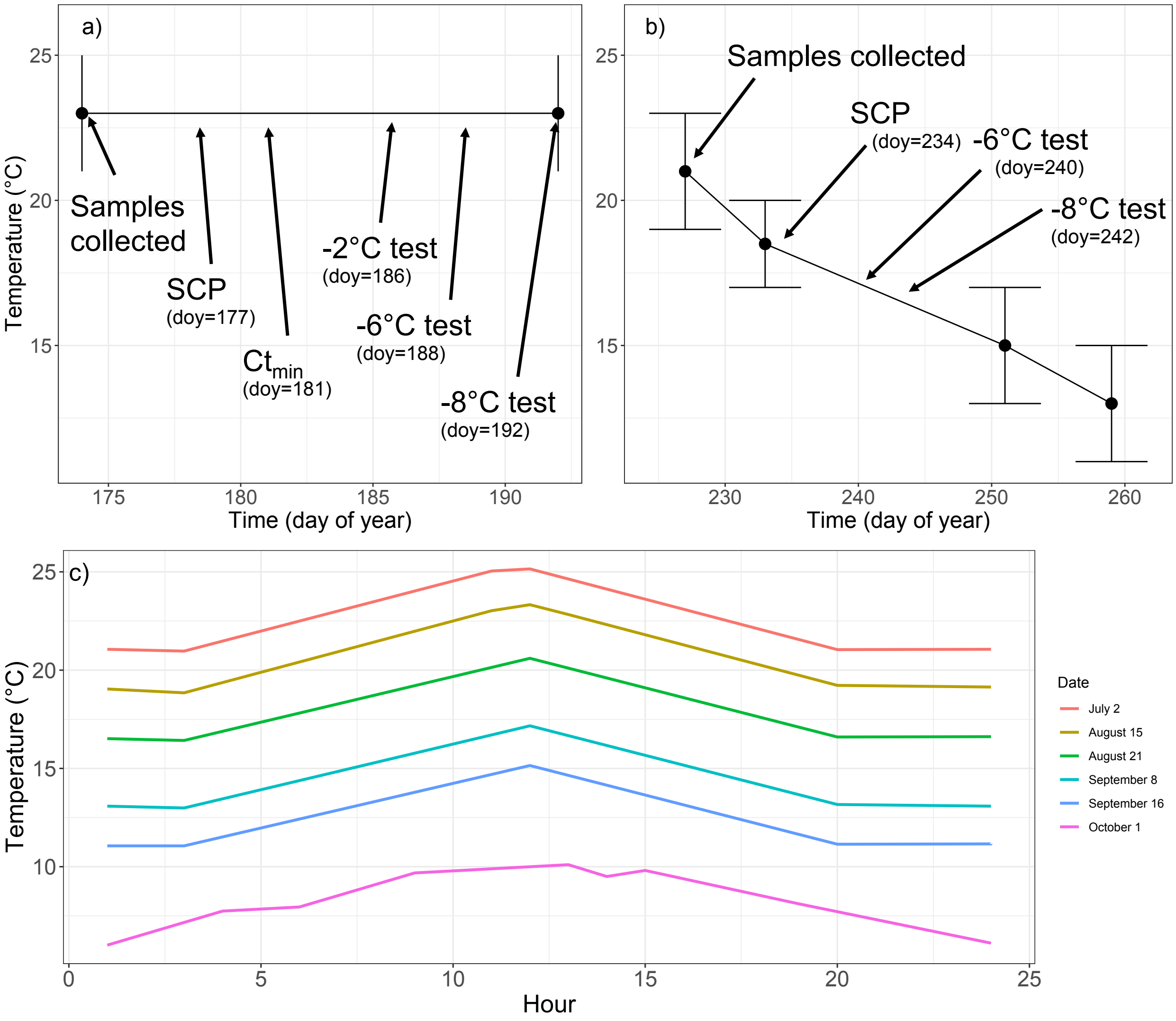
